# Seeing from the inside out: Intrinsic neural dynamics explain individual differences in movie watching

**DOI:** 10.64898/2025.12.03.692223

**Authors:** Masakazu Inoue, Masafumi Oizumi, Shuntaro Sasai

## Abstract

When viewing the same movie, individuals attend to different visual elements and construct distinct narratives. We hypothesized that these differences arise partly from intrinsic neural dynamics that specify upcoming visual content before eye movements occur, particularly when external visual saliency is low. Using fMRI and eye-tracking during a two-hour movie (n=12), we examined spontaneous gaze shifts during low versus high visual saliency periods. During low-saliency periods, when inter-individual variability was greatest, we found that visual features of upcoming gaze targets 2 seconds before eye movements could be predicted by right temporo-parieto-occipital junction (R-TPOJ) activity. Critically, the strength of connectivity between R-TPOJ and the right superior parietal lobule, which predicts gaze position, correlated with gaze-shift frequency during low-saliency periods. These findings suggest that R-TPOJ specifies the next scene content prior to foveation, and that eye movements serve to align this internally specified content with external stimuli, thereby shaping individual differences during movie-watching.

## Main

Humans actively experience the world in remarkably diverse ways. Even when watching the same movie, viewers do not uniformly attend to the same scenes, characters, or objects, nor do they form identical impressions^1–4^. Rather, the scenes that remain memorable, the characters that stand out, and the narratives constructed show substantial individual variation^5^. From the traditional perspective that the brain passively responds to information from the external environment^6–9^, such diversity is difficult to explain: if external input is identical, why do viewers engage in distinct patterns of visual exploration—seeking different information and directing their gaze toward different elements of the same scenes?

The answer may lie in reconsidering the primary causes of brain activity. The brain is a vast network of neurons, where cortical neurons largely fire in response to inputs from numerous other cortical and subcortical neurons projecting to them^10^. Crucially, these projecting neurons themselves receive input predominantly from other cortical neurons^11–13^, with only a small fraction receiving direct input from sensory organs such as the eyes and ears. This anatomical organization suggests that neural activity at any given moment is driven primarily by the brain’s own prior activity state.

This perspective^14,15^ contrasts with the traditional view^16,17^ that external stimuli (extrinsic input) are the primary determinants of brain activity. While certain cortical regions do show activation relevant to specific stimuli, recent research—including challenges such as the Algonauts Challenge^18^—demonstrates that external stimuli alone can explain only a portion of neural activity^19^; complete prediction remains difficult despite advances in neural network models^20^. This limited explanatory power suggests that understanding brain dynamics requires considering intrinsic dynamics: neural activity generated autonomously within the brain. Behavior and experience, therefore, are not simply reactions to environmental input but are shaped by the brain actively determining its own states.

Extending this perspective to naturalistic movie viewing, viewers may construct personal interpretations based on preceding scenes. Such intrinsically formed interpretations could constrain expectations about upcoming content and guide exploratory eye movements toward anticipated meaningful elements. Thus, individual differences in intrinsically constructed narratives should manifest as differences in visual exploration patterns during movie viewing.

We therefore hypothesized that individuals maintain internal visual features, and that active information sampling guided by these features generates patterns of visual exploration that are not accounted for by external stimuli alone (Fig. 1a). Under this assumption, gaze shifts occurring during low-saliency conditions should be more strongly shaped by intrinsic neural dynamics than by external sensory changes. If this hypothesis is correct, we predict: (1) brain regions exist whose activity predicts the visual semantic content at future gaze targets before eye movements occur, specifically during small saliency changes; and (2) brain regions exist that are involved in determining gaze destinations under small saliency change conditions.

**Fig. 1:**
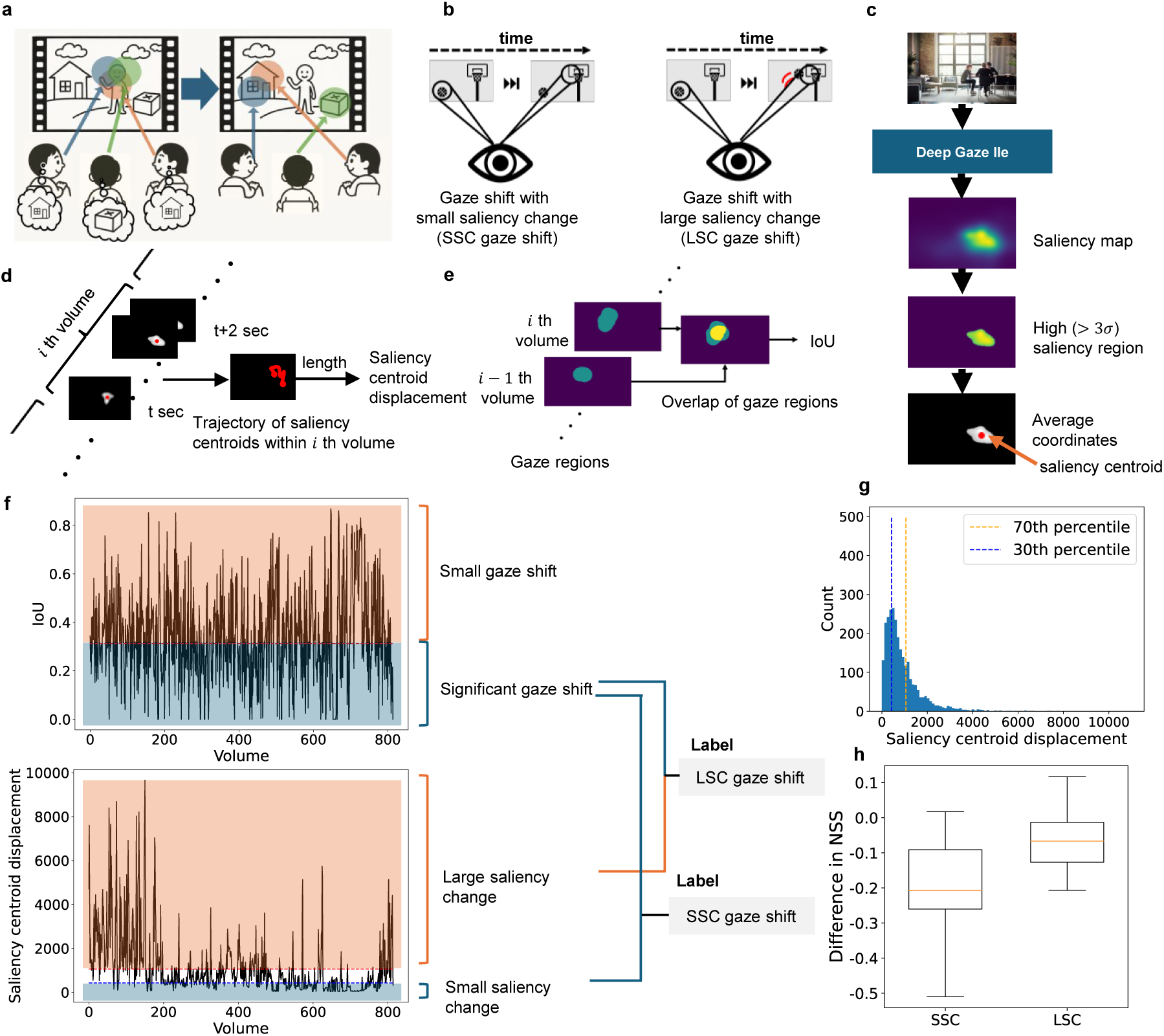
Classification of gaze shifts. **a**, Conceptual hypothesis: Individual differences in visual exploration are explained by intrinsic brain dynamics. **b**, Definition of gaze-shift labels: gaze shifts accompanied by large saliency changes (LSC gaze shift) and those with small saliency changes (SSC gaze shift). **c**, Using Deep Gaze IIE^23^, we generated saliency maps and calculated the average coordinates of high saliency regions (saliency centroids). **d**, The saliency centroid displacement was defined as the sum of squared Euclidean distances between centroids of consecutive frames within each 2-s window. **e**, The Intersection over Union (IoU) between gaze regions of the current and previous 2-s windows was computed to quantify the degree of gaze shift. **f**, Volumes whose IoUs were below the median (upper blue area) and whose saliency centroid displacements were below or above the 30th/70th percentile (lower blue/orange area) were labeled as SSC or LSC gaze shifts, respectively. **g**, The histogram of saliency centroid displacement shows that the 30th percentile approximately corresponds to its mode. **h**, Boxplots illustrating the participant-wise differences in Normalized Scanpath Saliency^24^ (NSS) between gaze-shift and gaze-fixation periods under the SSC and LSC conditions.

To test these predictions, we analyzed publicly available fMRI and eye-tracking data acquired simultaneously while participants watched a two-hour movie. Using these data, we identified the brain regions that support these predictive functions and examined whether coordinated activity between them accounts for individual differences in gaze behavior— specifically, the frequency of gaze shifts—under low-saliency conditions.

## Results

### Classification of gaze shifts

We prepared the data for analysis by processing and synchronizing multimodal time-series recordings. We utilized data from the StudyForrest project^21^, which simultaneously recorded 3T fMRI and eye-tracking data while participants viewed a two-hour movie. We analyzed 12 participants whose data showed no acquisition abnormalities. We synchronized video (25 FPS), eye-tracking (1000 Hz), and fMRI data (TR=2 seconds) by computing representative values at 2-second intervals. Eye-tracking data were classified into gaze events (fixations, saccades, post-saccadic oscillations, and pursuit) using REMoDNaV^22^. We focused on fixations and pursuit events, as objects targeted by these gaze events are perceived with relative stability.

Our framework relies on the hypothesis that individuals maintain internal visual features, and that active information sampling driven by these features generates patterns of visual exploration not accounted for by variations in external sensory input alone (Fig. 1a). Based on this hypothesis, we operationally defined two types of gaze shifts according to the magnitude of visual saliency change accompanying the shift (Fig. 1b): small saliency change (SSC) and large saliency change (LSC). To quantify saliency changes, we used DeepGaze IIE^23^ to generate saliency maps for each movie frame (Fig. 1c), calculated the average coordinates of regions with high saliency (saliency centroid), and measured the displacement of the centroid over each 2-second interval (Fig. 1d). In parallel, we computed the Intersection over Union (IoU) between consecutive gaze regions to quantify the extent of gaze shifts (Fig. 1e).

Based on these metrics, we labeled time points using a dual-criterion approach shown in Figure 1f: IoU values (to identify gaze shifts) and saliency centroid displacements (to characterize changes in extrinsic stimuli). Gaze shifts during small visual saliency changes (SSC gaze shifts) were identified when gaze shifted significantly (IoU below the median) despite minimal change in visual saliency (saliency centroid displacement below the 30th percentile, lower blue area in Fig. 1f). Conversely, gaze shifts during large visual saliency changes (LSC gaze shifts) occurred when significant gaze shifts (IoU below the median) coincided with large changes in visual saliency (saliency centroid displacement above the 70th percentile, lower orange area in Fig. 1f). We determined the thresholds for saliency centroid displacement based on the distribution, where the threshold for SSC gaze shifts corresponds to the mode (Fig. 1g). Examples of SSC/LSC gaze shifts are shown in Figure S1.

### Reduced inter-participant consistency during small saliency changes

Based on the event-labeled data derived from the procedures above, we found that gaze shifts were less consistent across participants under small saliency change (SSC) conditions, indicating greater variability in gaze behavior. We quantified inter-participant consistency in gaze behavior using the Normalized Scanpath Saliency (NSS) metric^24^. NSS measures the mean normalized saliency value at each observer’s gaze positions, with positive scores indicating gaze positions that align with those of other participants. Figure 1h shows the difference in mean NSS between gaze-shift and gaze-fixation samples (gaze-shift - fixation) for the SSC and LSC conditions. The SSC condition exhibited a larger negative NSS difference compared to the LSC condition (two-sided t-test, *p* < 0.1), indicating greater spatial dispersion of gaze positions following gaze shifts. This suggests that gaze shifts during SSC are more strongly influenced by environment-independent mechanisms than those during LSC.

### Visual features at future gaze targets can be decoded from R-TPOJ activity

To test our hypothesis that specific brain regions exist that actively specify the visual content at future gaze targets, we applied a decoding framework that identified ROIs whose activity predicted forthcoming gaze-region features. As illustrated in Figure 2a, each participant’s data was divided into training and test segments, and the cortex was parcellated into 44 ROIs^25^. Visual features at the gaze location were extracted using a pretrained Vision Transformer (ViT)^26,27^, and BOLD signals were time-shifted by Δτ (−6, −4, −2, 0, 2, 4, 6, 8, 10) seconds ahead to evaluate how early each ROI encoded this information. Building on prior work^28^, we trained linear decoders on the training dataset to predict low-dimensional visual-feature components from BOLD activity. Decoding accuracy was quantified in the test dataset using Pearson correlation coefficients for SSC and LSC events separately, yielding correlation scores for each participant, ROI, and time lag.

**Fig. 2:**
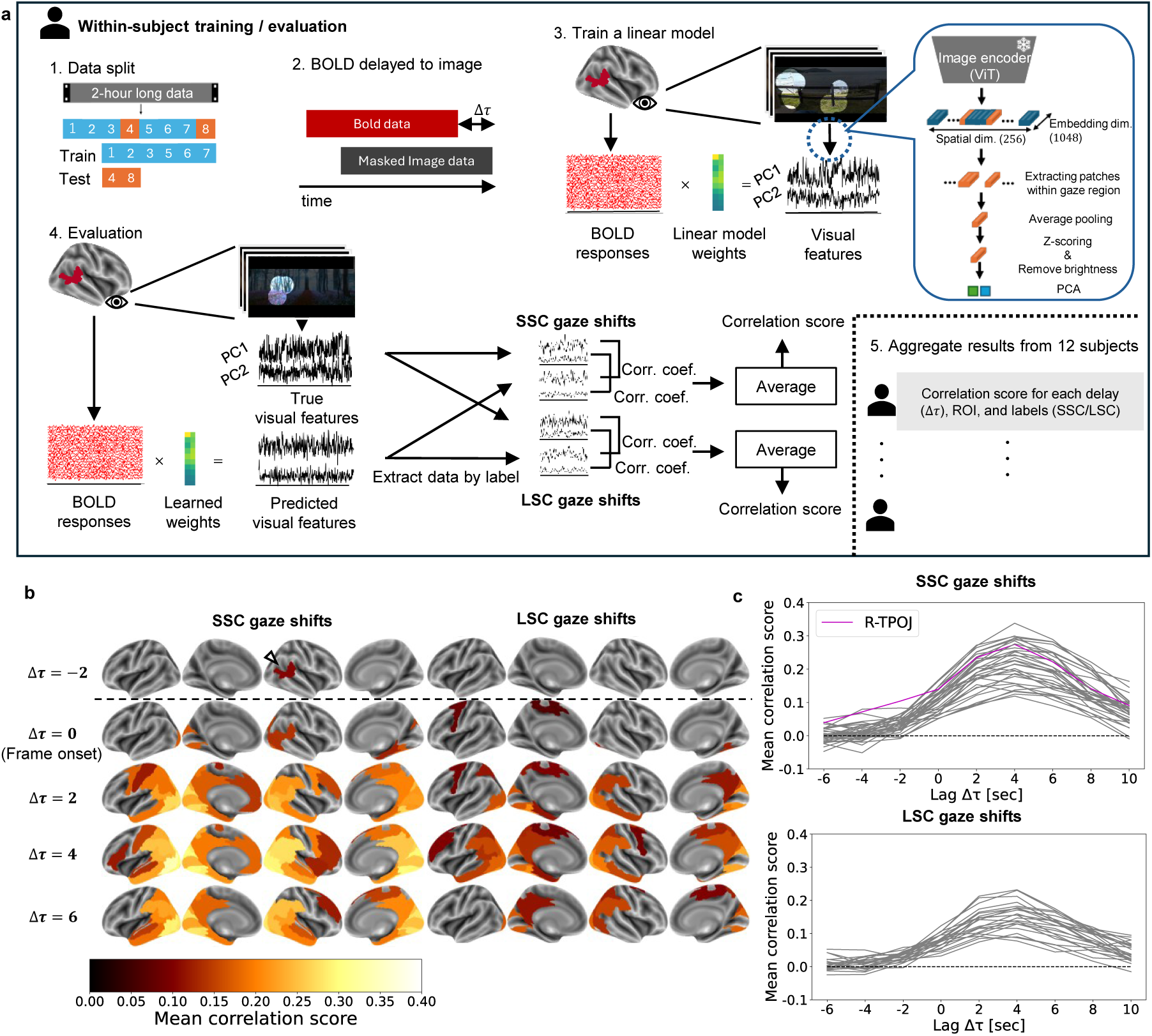
Decoding visual features from BOLD responses. **a**, Schematic of the decoding procedure. For each temporal lag Δτ (–6, –4, –2, 0, 2, 4, 6, 10 s), a linear model (ridge regression) was trained to predict the principal components of visual features of images masked based on gaze region, extracted by a Vision Transformer (ViT), from BOLD responses delayed by Δτ relative to image. Decoding performance was quantified as the correlation score, defined as the mean correlation coefficient between predicted and true component values and was assessed for test data separately for SSC or LSC gaze shifts. Model training and evaluation were repeated for each participant (N = 12), lag, and ROI. **b**, Spatiotemporal distribution of ROIs significantly predicting the visual features of gaze regions. ROIs with significant correlation scores (one-sided t-test, N = 12, Bonferroni-corrected, P < 0.05) at each Δτ are color-coded according to the mean correlation score across participants. The triangle marks the ROI whose correlation score became significant at negative lags (Δτ < 0) in the small-saliency-change (SSC) gaze-shift condition. **c**, Temporal profiles of correlation scores. Correlation scores across Δτ are shown for ROIs exhibiting significance at any lag. Colored traces indicate ROIs showing significant correlations at negative lags (Δτ < 0).

We identified the earliest brain regions capable of decoding visual features at gaze locations and found that only SSC conditions contained ROIs that predicted upcoming gaze-region features before gaze shifts occurred (Δτ<0). Figure 2b maps the spatial-temporal patterns of ROIs that significantly predicted visual features of the gaze region. Each panel shows results for different time lags (Δτ), with only significant ROIs colored (*t*-test, *N*=12, Bonferroni correction, α=0.05). Color intensity indicates group-mean correlation scores. For SSC gaze shifts, the Right Temporo-Parieto-Occipital Junction (R-TPOJ) was the earliest and unique ROI able to significantly predict visual features of the gaze region at Δτ=−2 (mean correlation=0.104, *p*<0.05, Bonferroni correction). In contrast, for LSC gaze shifts, no ROIs showed significant predictions at Δτ<0. There is no significant difference in auto correlation of visual features between SSC and LSC conditions (Fig. S3a, left panel), supporting that similarity of extrinsic stimuli across time cannot explain the difference in the decodability. We confirmed that R-TPOJ’s correlation score at Δτ=−2 is higher than that of other ROIs in the SSC condition, while no ROI showed these characteristics in the LSC condition (Fig. 2c). Comprehensive spatio-temporal pattern maps for time lags from Δτ=−6 to Δτ=10 are shown in Figure S2.

To assess whether prediction errors might arise not only from saliency change but also from factors such as gaze-shift magnitude or image-feature distributions, we examined the relationship between R-TPOJ’s predictive power and saliency change, image features, and the IoU between gaze regions, without splitting the data into SSC and LSC. Figure S3b shows the time series of prediction errors when predicting visual features at gaze positions from R-TPOJ activity at Δτ=-2. We found a weak correlation between these prediction errors and the magnitude of saliency change (Fig. S3c, corr.=0.1, *p*<0.01). This indicates that R-TPOJ’s influence on gaze control becomes stronger when the influence of external environmental input that triggers gaze shifts is weaker. Additionally, we found no relationship between image features or the IoU we used in our analysis (IoU<0.3) and R-TPOJ’s prediction error (Fig. S3d-e).

### Visual features outside future gaze targets cannot be decoded from any ROI

To test whether R-TPOJ activity selectively encodes visual features of the actual gaze targets—or whether features outside the gaze regions could also be decoded—we performed control evaluations under two alternative target locations. Under saliency-mask conditions, we evaluated predictions using visual features from the most salient regions rather than actual gaze regions (Fig. 3a). Under shuffled-mask conditions, we used temporally permuted actual gaze regions (Fig. 3b). Neither control condition yielded significant correlations at Δτ=−2 for either event type, confirming that our models specifically predicted visual features of future gaze regions rather than salient or random image regions (Fig. 3c–f). Comprehensive spatio-temporal pattern maps for time lags from Δτ=−6 to Δτ=10 are shown in Figures S4–5.

**Fig. 3:**
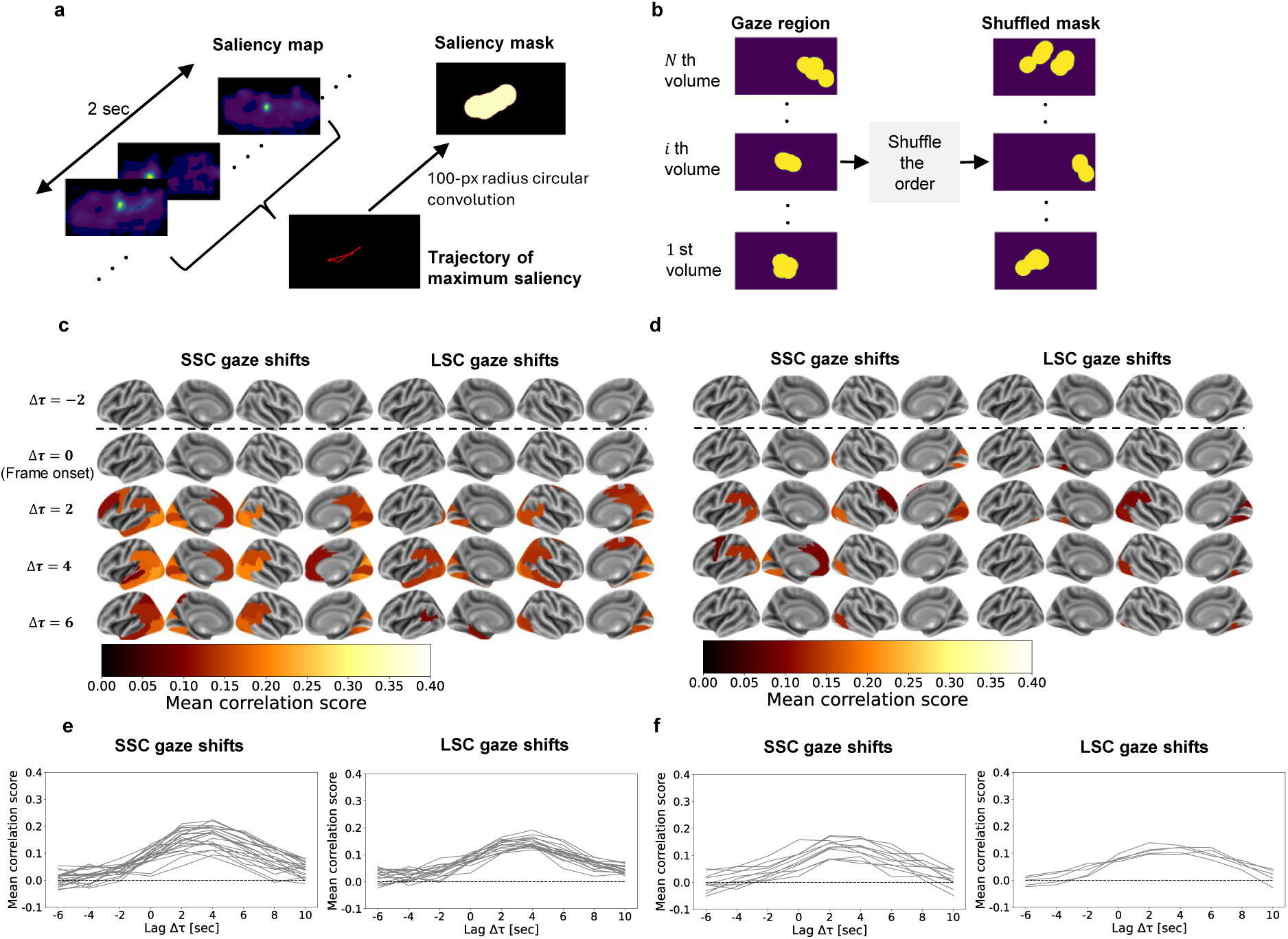
Decoding visual features outside the gaze region. **a**, Construction of the saliency mask. Within each 2-s window, regions of high saliency were retained by convolving a circular kernel with a radius of 100 pixels along the trajectory of maximum saliency values in the saliency map and blacking out all other regions. **b**, Construction of the shuffled mask. Masks indicating gaze regions were temporally shuffled by randomly permuting their order, thereby generating masks that differed from the actual ones. **c**, Spatiotemporal distribution of ROIs significantly predicting the visual features of saliency-masked images. ROIs with significant correlation scores (one-sided t-test, N = 12, Bonferroni-corrected, P < 0.05) at each Δτ are color-coded according to the mean correlation score across participants. **d**, Spatiotemporal distribution of ROIs significantly predicting the visual features of shuffled-masked images. ROIs with significant correlation scores (one-sided t-test, N = 12, Bonferroni-corrected, P < 0.05) at each Δτ are color-coded according to the mean correlation score across participants. **e**, Temporal profiles of correlation scores under the saliency-mask condition. Correlation scores across Δτ are shown for ROIs exhibiting significance at any lag. **f**, Temporal profiles of correlation scores under the shuffled mask condition. Correlation scores across Δτ are shown for ROIs exhibiting significance at any lag.

These findings demonstrate that R-TPOJ’s activity is specific to visual features of unseen gaze targets during SSC gaze shifts.

### Future gaze positions cannot be predicted from R-TPOJ activity

Our previous analyses revealed that R-TPOJ activity can decode the content of future gaze targets. Next, we examined the relationship between R-TPOJ and the brain mechanisms underlying gaze shifts themselves. Brain regions involved in gaze shifts should be able to estimate the spatial location toward which gaze is directed. Therefore, we sought to identify brain regions from which the spatial location of future gaze targets could be decoded.

Figure 4a illustrates our decoding approach for gaze positions. Gaze position was defined as the average coordinates of fixation and pursuit endpoints within each 2-second interval. Similar to the visual feature analysis, we trained linear models via Ridge regression to decode the *x* and *y* coordinates of gaze positions from BOLD signals in the training dataset. For evaluation, we extracted SSC and LSC events separately in the test dataset, predicted coordinates from BOLD signals for each event type, calculated Pearson correlations between predicted and true coordinates, and averaged them to obtain correlation scores. This procedure yielded correlation scores for each participant, ROI and time lag. We excluded L/R-Orbital and Polar Frontal areas due to eye movement artifacts in these regions^29,30^.

**Fig. 4:**
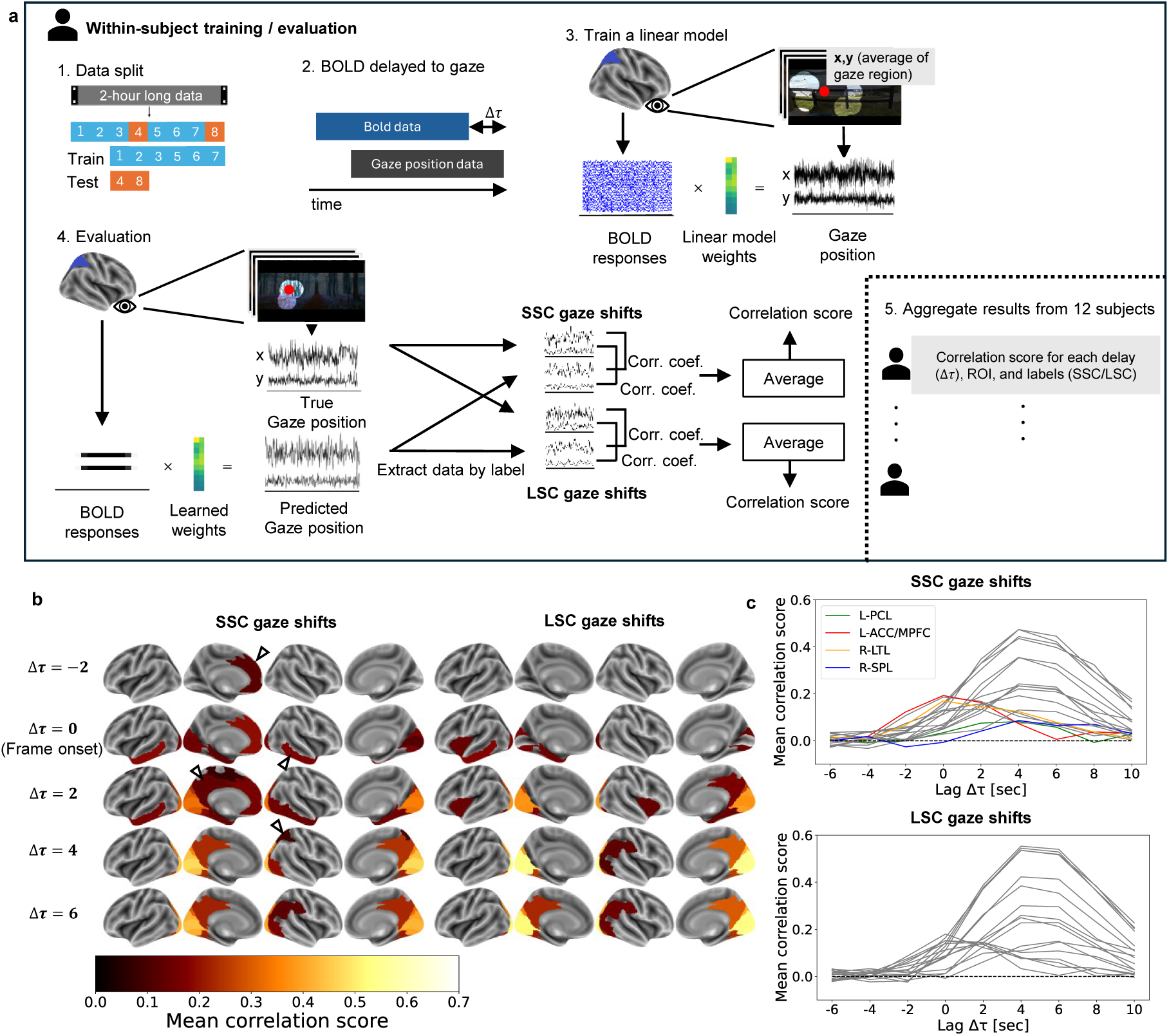
Decoding gaze positions from BOLD responses. **a**, Schematic of the decoding procedure. For each temporal lag Δτ (–6, –4, –2, 0, 2, 4, 6, 10 s), a linear model (ridge regression) was trained to predict the x and y coordinates of gaze positions, averaged gaze region over a 2-s window, from BOLD responses delayed by Δτ relative to gaze positions. Decoding performance was quantified as the correlation score, defined as the mean correlation coefficient between the predicted and true coordinates and was assessed for test data separately for SSC or LSC gaze shifts. Model training and evaluation were repeated for each participant (N = 12), lag, and ROI. **b**, Spatiotemporal distribution of ROIs significantly predicting gaze positions. ROIs with significant correlation scores (one-sided t-test, N = 12, Bonferroni-corrected, P < 0.05) at each Δτ are color-coded according to the mean correlation score across participants. The triangles mark the ROIs that showed significance specifically in the small-saliency-change (SSC) gaze-shift condition. **c**, Temporal profiles of correlation scores. Correlation scores across Δτ are shown for ROIs exhibiting significance at any lag. Colored traces indicate ROIs showing significant correlations for SSC gaze shifts only.

Based on the decoding approach, we identified a set of brain regions that significantly predicted gaze positions specifically during SSC conditions across time lags from Δτ = −6 to Δτ = 10. Figure 4b maps the spatial-temporal patterns of ROIs significantly predicting gaze positions. Each panel displays results for different time lags (Δτ), with only significant ROIs colored (*t*-test, *N*=12, Bonferroni correction, *α*=0.05). Color intensity indicates group-mean correlation scores. For both gaze shift conditions (Fig. 4b and 4c), a largely overlapping set of regions significantly estimated gaze position, including visual cortices, temporal areas, and some parietal and cingulate regions. However, careful comparison of the two rows reveals that the LSC pattern did not include L-Anterior Cingulate and Medial Prefrontal cortex (L-ACC/MPFC), L-Paracentral lobular and Mid Cingulate (L-PCL), R-Lateral Temporal lobe (R-LTL), and R-Superior Parietal lobule (R-SPL), distinguishing it from the pattern in the SSC condition. This dissociation indicates that these regions are specifically involved in gaze shifts that occur independently of saliency changes. Notably, R-TPOJ activity did not significantly predict gaze position under either condition, indicating that the neural system determining the location of gaze shifts is distinct from the system specifying the visual content of upcoming gaze targets. Comprehensive spatio-temporal pattern maps for time lags from Δτ=−6 to Δτ=10 are shown in Figure S6.

### R-TPOJ’s functional connectivity explains individual differences in SSC gaze shifts

Thus far, we confirmed that SSC events, in which environmental input triggering gaze shifts is minimal, show greater inter-individual variance in gaze positions following gaze shifts. We also found that only during SSC events does a region exist (R-TPOJ) that can predict the visual features at future gaze targets. Furthermore, this region cannot specify “where to look,” indicating that determining gaze content and determining gaze location are mediated by distinct systems. These findings raise the question of whether the degree of functional coupling between the region harboring content representations (R-TPOJ) and the regions involved in determining gaze locations under SSC conditions contributes to individual differences in environment-independent gaze-shift frequency.

To evaluate the relationship between functional coupling and gaze-shift behavior, we quantified functional connectivity (FC) and behavioral indices using the following procedure. Figure 5a illustrates our functional connectivity calculation method. We calculated FC between R-TPOJ as the seed ROI and each of the four ROIs (L-ACC/MPFC, L-PCL, R-LTL, and R-SPL) predicting gaze positions as targets by taking the mean absolute correlation coefficient (MACC) across voxel pairs. Each MACC was then normalized by a baseline defined as the average connectivity between R-TPOJ and all remaining ROIs, allowing us to assess region-specific coupling during the task, independent of overall brain connectivity^31–33^. To relate these FC measures to behavior, we derived two indices capturing individual gaze-shift tendencies: the probability of SSC gaze shifts (number of SSC gaze shifts / number of SSC events) and the probability of LSC gaze shifts (number of LSC gaze shifts / number of LSC events), reflecting how often each participant shifted gaze under small versus large saliency-change conditions

**Fig. 5:**
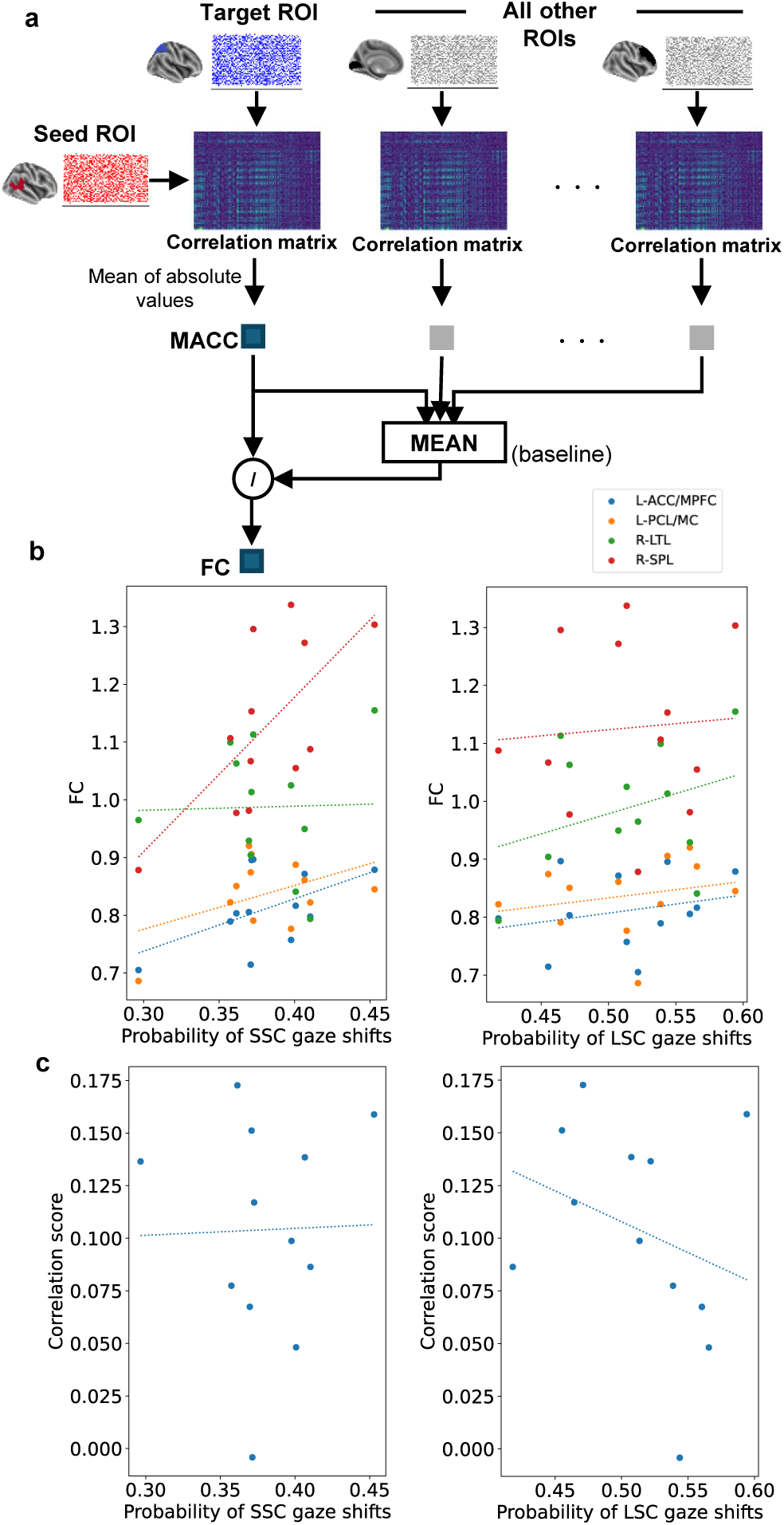
Correlation analysis between functional connectivity and gaze shift probability. **a**, Computation of functional connectivity (FC). To obtain FC between the seed ROI and a target ROI, the mean absolute correlation coefficient (MACC) was computed between voxels in the seed and target ROIs across samples in the test dataset, and divided by the baseline, defined as the mean MACC between the seed ROI and all other ROIs. **b**, Scatter plots showing the relationship between FC and the probability of SSC (left) and LSC (right) gaze shifts across participants. Each dot represents one participant. Dashed lines denote the fitted regression lines. Colors indicate target ROIs of FCs. The seed ROI was the right temporo-parieto-occipital junction (R-TPOJ). **c**, Scatter plots showing the relationship between correlation scores obtained from decoding visual features of gaze-region-masked images from the BOLD response of the R-TPOJ at 2 s before gaze shifts, and the probability of SSC (left) and LSC (right) gaze shifts. Dashed lines denote the fitted regression lines.

We identified functional connectivity that correlated with individual differences in gaze-shift frequency. Figure 5b presents scatter plots of FC values versus the probability of SSC gaze shifts (number of SSC gaze shifts / number of SSC events) or LSC gaze shifts for each subject (N = 12). As shown in Figure 5b, we found that functional connectivity between R-TPOJ and R-SPL significantly correlated with SSC gaze shift probability (*r*=0.684, surrogate test: 1,000 permutations × 10 seeds; mean rank=7.5, *p*<0.05). Participants with stronger coupling between these regions showed a higher tendency to shift their gaze during low saliency changes. The other three FC measures (L-Anterior Cingulate and Medial Prefrontal cortex, L-Paracentral lobular and Mid Cingulate, and R-Lateral Temporal lobe) did not show significant correlations with SSC probability. Figure 5c displays analogous scatter plots for FC values versus the probability of LSC gaze shifts. We found that FC between R-TPOJ and L-ACC/MPFC shows marginally significant anti-correlation (*r*=-0.676, surrogate test: 1,000 permutations × 10 seeds; mean rank=987.5, *p*<0.1) with partial correlation analysis, indicating that stronger coupling between these regions prevents the occurrence of gaze shifts triggered by saliency changes.

To validate the analysis results in Figure 5b, we constructed a model to predict each participant’s gaze shift probability from functional connectivity and performed verification using the model’s error through a leave-one-subject-out approach (Fig. S7a). As a result, we confirmed that even with this method, the functional connectivity between R-TPOJ and RSPL best predicted the gaze shift probability of test subjects in the SSC condition (Fig. S7b). These findings demonstrate that functional interaction between a system for intrinsically specifying next scene content and that for gaze positioning shapes individual tendencies in viewing behavior.

## Discussion

Our study demonstrates that brain activity can specify visual features at future gaze targets before eye movements occur, specifically when visual saliency changes are minimal. Using simultaneous fMRI and eye-tracking during movie viewing, we found that activity in the right temporo-parieto-occipital junction (R-TPOJ) enabled decoding of visual features at upcoming gaze regions 2 seconds before gaze shifts, but only during small saliency change (SSC) conditions—when external environmental factors triggering gaze shifts were weak. Critically, R-TPOJ activity did not enable decoding of visual features at non-gaze locations, nor could it be used to decode the spatial coordinates of future gaze targets. This dissociation suggests that gaze shift planning involves neural systems distinct from R-TPOJ.

Beyond these core findings that R-TPOJ encodes the visual content to be seen at future gaze targets, we identified specific brain regions (L-Anterior Cingulate and Medial Prefrontal cortex, L-Paracentral lobular and Mid Cingulate, R-Lateral Temporal lobe, and R-Superior Parietal lobule) that are selectively recruited during SSC gaze shifts and encode the gaze destination. Importantly, functional connectivity between R-TPOJ and R-Superior Parietal lobule positively correlated with gaze shift frequency during SSC conditions, while connectivity between R-TPOJ and L-Anterior Cingulate and Medial Prefrontal cortex negatively correlated with gaze shifts during LSC conditions. These findings suggest that individual differences in movie-watching behavior emerge from intrinsic neural dynamics that actively specify visual content to be explored.

To clarify the temporal characteristics and specificity of the information that R-TPOJ represents about future gaze targets, we next detail the temporal profile and specificity of the visual information represented in this region. Our results confirmed that the R-TPOJ carries visual information specific to future gaze-target regions rather than non-gaze regions. The optimal time shift (Δτ) at which BOLD signals from most ROIs best enabled decoding of visual features was +4 seconds, consistent with the hemodynamic response function^34,35^ and reflecting neural responses to visual input at the gaze location. R-TPOJ also showed peak decoding performance at Δτ = +4 seconds, but critically, decoding from R-TPOJ was possible at Δτ = −2 seconds—before gaze shifts occurred—specifically during SSC conditions (Fig. 2b). This early capability indicates that R-TPOJ activity is not reactive to visual input but reflects its intrinsic specification of upcoming visual content. The specificity of this decoding was further validated through control analyses using saliency-based and temporally shuffled masks. Models trained on each mask condition failed to show significant correlations under these control conditions (Fig. 3), demonstrating that decoding from R-TPOJ activity specifically reflected content at future gaze targets during SSC events rather than capturing information from salient or arbitrary spatial locations.

The functional characteristics we observed in R-TPOJ align with its known anatomical and functional properties. R-TPOJ has been implicated in distinguishing self from others^36^ and in shifting attention and belief states^37^ —functions requiring comparison of intrinsic information with external inputs. Our finding that R-TPOJ activity relates to visual features to be seen in the near future is consistent with this comparative role. Anatomically, TPOJ is located at the posterior end of the Sylvian fissure^38^, where the temporal, parietal, and occipital lobes converge, and has been proposed as a hub linking local and distant multisensory cortical regions through extensive white matter connections. This anatomical connectivity suggests TPOJ is ideally positioned to integrate multimodal sensory expectations with current activity states across distributed cortical areas, thereby specifying the next brain state to be realized through sensory sampling.

In contrast to the R-TPOJ, which represents visual features of future gaze targets, several other regions—L-Anterior Cingulate and Medial Prefrontal cortex (L-ACC/MPFC), L-Paracentral lobular and Mid Cingulate (L-PCL), R-Lateral Temporal lobe (R-LTL), and R-Superior Parietal lobule (R-SPL)—encoded the gaze position during SSC events. These regions align well with known components of the voluntary oculomotor network^39–41^. ACC/MPFC is functionally connected with frontal eye fields^42^ and involved in voluntary gaze shifts following internal goals^43^. R-SPL is extensively connected with the oculomotor network^39,44,45^ and integrates somatosensory input^46^ with visual information to guide spatial attention^47^ and coordinate gaze movements^48^. Their selective recruitment during SSC conditions suggests they specifically support gaze shifts driven by intrinsic dynamics rather than by salient external stimuli.

Because eye movements can confound fMRI measurements near orbital regions, we carefully examined and eliminated these artifacts to ensure that our decoding reflected neural—not artifactual—activity. Eye movements and extraocular muscle contractions induce magnetic field changes that affect fMRI signals in adjacent orbital frontal regions^29,30^, producing correlation peaks at Δτ = 0. Indeed, ROIs showing such early peaks were predominantly located in frontal regions near the eyes, confirming these reflect artifacts rather than genuine gaze planning signals. By excluding these regions, we ensured that our identified ROIs reflect neural activities involved in gaze control.

Our functional connectivity analysis further suggests that exploratory gaze shifts arise from coordinated interactions between regions that encode visual features of future gaze locations, such as the R-TPOJ, and regions that support gaze control, such as the R-SPL. Functional connectivity between R-SPL and R-TPOJ was positively correlated with gaze shift probability during SSC conditions (Fig. 5b). Participants showing stronger coupling between these regions exhibited higher gaze shift frequencies during periods of low external saliency, whereas this connectivity did not explain gaze shifts during LSC conditions. This finding demonstrates that R-TPOJ (from which visual content at future gaze locations can be decoded) and R-SPL (from which gaze destinations can be decoded) work in concert to drive voluntary exploratory eye movements. The coupling between these systems appears critical for individual differences in exploratory viewing behavior when external guidance is minimal.

These results collectively indicate that intrinsic neural dynamics dominate gaze behavior when external stimulus-driven guidance is weak. Traditional models of gaze control have attributed eye movements primarily to environmental inputs such as saliency changes^6–9^, yet gaze shifts still occur with substantial individual variability even in the absence of large saliency fluctuations (Fig. 1h). Our findings suggest that under such conditions, intrinsic neural dynamics—activity patterns generated autonomously by ongoing brain activity— become the dominant driver. The ability to decode visual features of future gaze locations specifically during SSC conditions suggests that exploratory gaze shifts are directed toward visual content already represented in the R-TPOJ when external guidance is weak. These findings are consistent with the idea that the brain integrates the unfolding narrative to anticipate semantic content likely to appear next. The functional connectivity between R-TPOJ and gaze planning regions (particularly R-SPL) explained individual differences in gaze shift frequency during SSC, indicating that the ease with which intrinsic dynamics drive gaze shifts depends on coordination between narrative construction and gaze planning systems. This framework reframes visual exploration not as a passive response to environmental features but as an active behavior whereby the brain seeks visual content matching its current intrinsic state.

Our study has limitations that should be acknowledged. First, the sample size was relatively small (N=12), which may limit generalizability. However, we employed rigorous machine learning approaches, splitting data into training and test sets and validating models on held-out data. This cross-validation ensured our findings reflect genuine relationships rather than overfitting. The successful generalization to unseen test data and consistent patterns across participants provide confidence in our results despite the limited sample size. Nevertheless, future studies with larger samples would be valuable.

Second, our study demonstrated that brain activity can specify future gaze behavior, but we did not establish causal relationships. The correlational nature of fMRI data means we cannot definitively determine whether R-TPOJ activity causally drives gaze shifts or reflects parallel processes preceding eye movements. Establishing causality would require perturbation experiments—for example, using transcranial magnetic stimulation^49^ to temporarily disrupt R-TPOJ function and observe effects on gaze behavior, or optogenetic manipulation^50^ in animal models to test whether disrupting analogous regions affects anticipatory gaze patterns during naturalistic visual exploration. Such causal interventions await future research.

Additional considerations include that our findings are based on movie-viewing behavior, which is a specific form of naturalistic vision. While movies provide rich, ecologically valid stimuli, they differ from real-world visual exploration (e.g., constrained to 2D screen space, no agency over camera viewpoint). Future work should examine whether similar intrinsic dynamics govern gaze control during free exploration of 3D environments. Furthermore, investigating longer-timescale narrative construction and its relation to viewing behavior could provide additional insights into temporal dynamics.

In conclusion, our findings demonstrate that individual differences in movie-watching behavior emerge not solely from differences in how brains respond to external stimuli, but from differences in intrinsic neural dynamics that actively specify visual content to be explored. Activity in the right temporo-parieto-occipital junction relates to visual features at future gaze locations before eye movements occur, specifically when external visual guidance is minimal. The strength of functional coupling between this region and brain areas involved in gaze positioning explains individual variation in exploratory viewing behavior. These results suggest that visual exploration during naturalistic viewing is best understood not as a passive reaction to salient features in the environment, but as an active behavior whereby the brain seeks visual input that aligns with its internally constructed narrative. This perspective fundamentally reframes our understanding of how humans experience visual media, highlighting the central role of intrinsic brain dynamics in shaping subjective experience.

## Methods

### Participants and dataset

We used publicly available fMRI and eye-tracking data collected simultaneously while participants viewed a two-hour movie (resolution: 1280 × 720 pixels) from the Study Forrest project^14^. We analyzed data from twelve participants showing no abnormalities in fMRI data acquisition. All participants provided informed consent, and the study was approved by the ethics committee of Otto-von-Guericke University Magdeburg. The movie, fMRI, and eye-tracking data were divided into eight segments. Segments 1, 2, 3, 5, 6, and 7 were used as the training dataset, while segments 4 and 8 were reserved for evaluation.

### fMRI data preprocessing

fMRI data (3T scanner, TR = 2s) were preprocessed using fMRIPrep^51^. Following Glasser’s approach^19^, we parcellated cortices into 44 ROIs based on the HCP-MMP1.0 atlas. For each ROI, we extracted the average BOLD signal across all voxels at each time point. Consequently, the number of volumes obtained for segments 1–8 was 451, 441, 438, 488, 462, 439, 542, and 338, respectively.

### Eye-tracking data processing

Eye-tracking data (1000 Hz) were processed using gaze events classified by REMoDNaV^22^. We extracted gaze events classified as fixations and pursuit movements since objects are perceived stably during these two gaze types^22,52^. Each event entry included the label, onset time, duration, start and end coordinates on the display, amplitude, and velocity.

### Data temporal alignment

To analyze movie data (25 FPS) and eye-tracking data (1000 Hz) with fMRI data (TR=2s), we computed representative values every 2 seconds. Movie data were downsampled to 0.5 FPS. Thus, the 𝑛 th frame corresponds to the start of the 𝑛 th slice timing in the fMRI signal. For each 2-second interval, we obtained gaze regions by convolving gaze event trajectories occurring during that interval with a circle of 100-pixel radius (corresponding to 1.85° visual angle). Gaze positions were obtained by averaging endpoint coordinates of gaze events within each 2-second interval.

### Saliency computation

We obtained saliency maps for each movie frame using Deep Gaze IIE^23^ and calculated high-saliency region displacement over each 2-second interval (Fig. 1f). Saliency maps are matrices of the same size as the input image, normalized using a softmax function. From each saliency map within a 2-second window, we extracted pixels with values exceeding three standard deviations above the mean and calculated the average coordinates of these pixels as the centroid of high saliency region. The sum of squared Euclidean distances between centroids of consecutive frames within each 2-second window was defined as the saliency centroid displacement (Fig. 1d). In parallel, we computed the Intersection over Union (IoU) between consecutive gaze regions to quantify the extent of gaze shifts (Fig. 1e).

### Labeling

Based on the computed saliency centroid displacements, we defined gaze shift types as follows. For each sample, we assigned labels based on saliency displacement and gaze region overlap. We labeled instances as ‘gaze shift with small saliency change’ (SSC gaze shifts) when saliency centroid displacement was below the 30th percentile of its distribution (Fig. 1f, lower blue area) and IoU between current and previous gaze regions was below its median (Fig. 1f, upper blue area). Conversely, we labeled instances as ‘gaze shift with large saliency change’ (LSC gaze shifts) when the saliency displacement exceeded the 70th percentile (Fig. 1f, lower orange area) and gaze region overlap was below the median (Fig. 1f, upper blue area). We determined the thresholds for saliency shift distance based on the distribution, where the threshold for SSC gaze shifts corresponds to its peak (Fig. 1g). To ensure balanced sample sizes between conditions, the threshold for LSC gaze shifts was set to 70th percentile. As a result, we obtained more than 100 samples per participant, on average, for both LSC and SSC gaze shift conditions in the test dataset.

### Normalized Scanpath Saliency (NSS)

To quantify the degree of consistency in gaze behavior across observers, we employed the Normalized Scanpath Saliency (NSS) metric^53^, extended to the temporal domain^24^. Gaze samples belonging to fixation and pursuit events were extracted using a sliding window of 250 ms with a step size of 125 ms. Within each window, the gaze samples of all participants except one were used to construct a spatiotemporal fixation map by superimposing three-dimensional Gaussian kernels centered at each participant’s gaze position and time point (𝑥, 𝑦, 𝑡). The resulting fixation map 𝐹(𝑥, 𝑦, 𝑡) was normalized to zero mean and unit standard deviation—estimated via Monte Carlo sampling (4,000 samples)—to obtain the NSS map 𝑁(𝑥, 𝑦, 𝑡). The NSS score for the left-out participant was computed as the mean of 𝑁(𝑥, 𝑦, 𝑡) values at their gaze positions, following a leave-one-out cross-validation procedure.

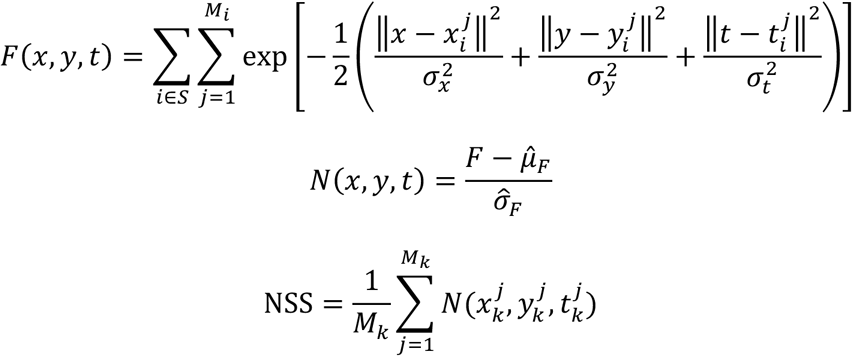

Here, 𝑀*_i_* denotes the number of gaze samples for participant 𝑖 within a window, and 𝑆 represents the set of all participants excluding the test subject 𝑘. Gaussian kernel parameters were set to 𝜎*_x_* = 𝜎*_y_* = 100 pixels (corresponding to 1.85° of visual angle) and 𝜎*_t_* = 160 ms (corresponding to a period of four video frames).

To match the sampling rate of the fMRI data, NSS values were aggregated and averaged every 2 seconds. Because the NSS value is normalized, random gaze distributions yield an expected score of zero, whereas positive scores indicate greater coherence of gaze behavior among observers.

### Prediction of visual features from brain activity

To identify ROIs that encode visual features of forthcoming gaze regions, we used BOLD signals temporally shifted relative to the visual features by Δτ seconds. Visual features were extracted from gaze-region–masked images using a pretrained Vision Transformer (ViT)^26^ from the BLIP Visual-Language Model^27^. For each 2-second interval, all image regions outside the gaze location were masked, and the masked image was passed through the ViT. Feature vectors were obtained from the final hidden layer by averaging patch embeddings within the gaze region. These features were Z-scored and orthogonalized with respect to global brightness (Fig. 2a). Principal component analysis (PCA) was then applied, and only the first two components were retained, based on pilot analyses indicating that higher components were poorly predicted from the BOLD signals.

Following the method of Horikawa et al.^28^, we trained linear models via Ridge regression^54,55^ using the 250 most correlated voxels from each ROI to predict the principal components of visual features from BOLD signals in the training dataset. Model performance was evaluated by applying trained models to the test dataset to obtain predicted component time series. Then, we extracted time points belonging to SSC and LSC events separately, predicted the principal components from BOLD signals for each event type, calculated Pearson correlation coefficients between predicted and true components, and averaged these correlations to obtain correlation scores. This procedure was repeated for each participant, principal component, Δ 𝜏 (−6, −4, −2, 0, 2, 4, 6, 8, 10 seconds), and ROI.

### Prediction of gaze positions from brain activity

To identify ROIs specifically involved in gaze shifts, we trained linear models predicting gaze positions from each ROI’s BOLD signals and examined ROIs specifically involved in predicting gaze positions during SSC gaze shifts. Utilizing BOLD signals time-shifted relative to the gaze positions by Δ𝜏 seconds, we constructed linear models as follows. First, we selected 250 voxels from each ROI whose BOLD signals showed the highest Pearson correlation^30^ with x or y coordinates. We then trained linear models by Ridge regression^54,55^. Trained models were applied to test data to obtain predicted coordinates (Fig. 4a). This procedure was repeated for each participant, coordinate, Δ𝜏 (−6, −4, −2, 0, 2, 4, 6, 8, 10 seconds), and ROI. We calculated correlation scores between predicted and true principal components for each gaze shift label. Left/right orbital and polar frontal regions were excluded due to known eye-movement-related BOLD artifacts^29,30^

### Functional connectivity analysis

To examine whether functional connectivity (FC) between the seed ROI predicting visual features at future gaze targets (R-TPOJ) and each target ROI predicting gaze positions (L-ACC/MPFC, L-PCL/MC, R-LTL, and R-SPL) could explain inter-subject differences in the probability of SSC/LSC gaze shifts, we calculated FC between seed and target ROI. For each pair, we computed the mean of absolute correlation coefficients (MACC) between all voxel pairs from the selected voxels in the seed and target ROIs across samples in the test dataset (Fig. 5a). To control for global connectivity effects during movie viewing, we normalized each MACC by dividing by the baseline, defined as the average MACC between R-TPOJ and all other ROIs. This normalization ensures that our FC measure reflects specific coupling between seed and target regions, independent of overall brain connectivity^31–33^ during the task.

We defined two behavioral measures to quantify individual differences in gaze shift tendencies: “probability of SSC gaze shifts” (number of SSC gaze shifts / number of SSC events) and “probability of LSC gaze shifts” (number of LSC gaze shifts / number of LSC events). These probabilities capture the tendency of each participant to shift their gaze under small versus large saliency change conditions.

We performed correlation analysis and partial correlation analysis to associate FC with probability of SSC gaze shifts and probability of LSC gaze shifts, controlling probability of gaze shifts.

### Statistical analysis

Statistical significance of mean correlation coefficients was assessed using t-tests (N=12, one-sided, significance level=0.05). Bonferroni correction was applied across ROIs, Δτ values, conditions (SSC/LSC), and mask types. For FC analyses, p-values were derived from rank-based permutation tests (1,000 shuffles, repeated 10 times with different random seeds). Mean ranks were converted to *p*-values and Bonferroni correction applied for the number of target ROIs (significance level=0.05). All analyses were performed using Python 3.8 with NumPy, SciPy, and scikit-learn libraries.

## Supplementary Figures

**Supplementary Fig. 1:**
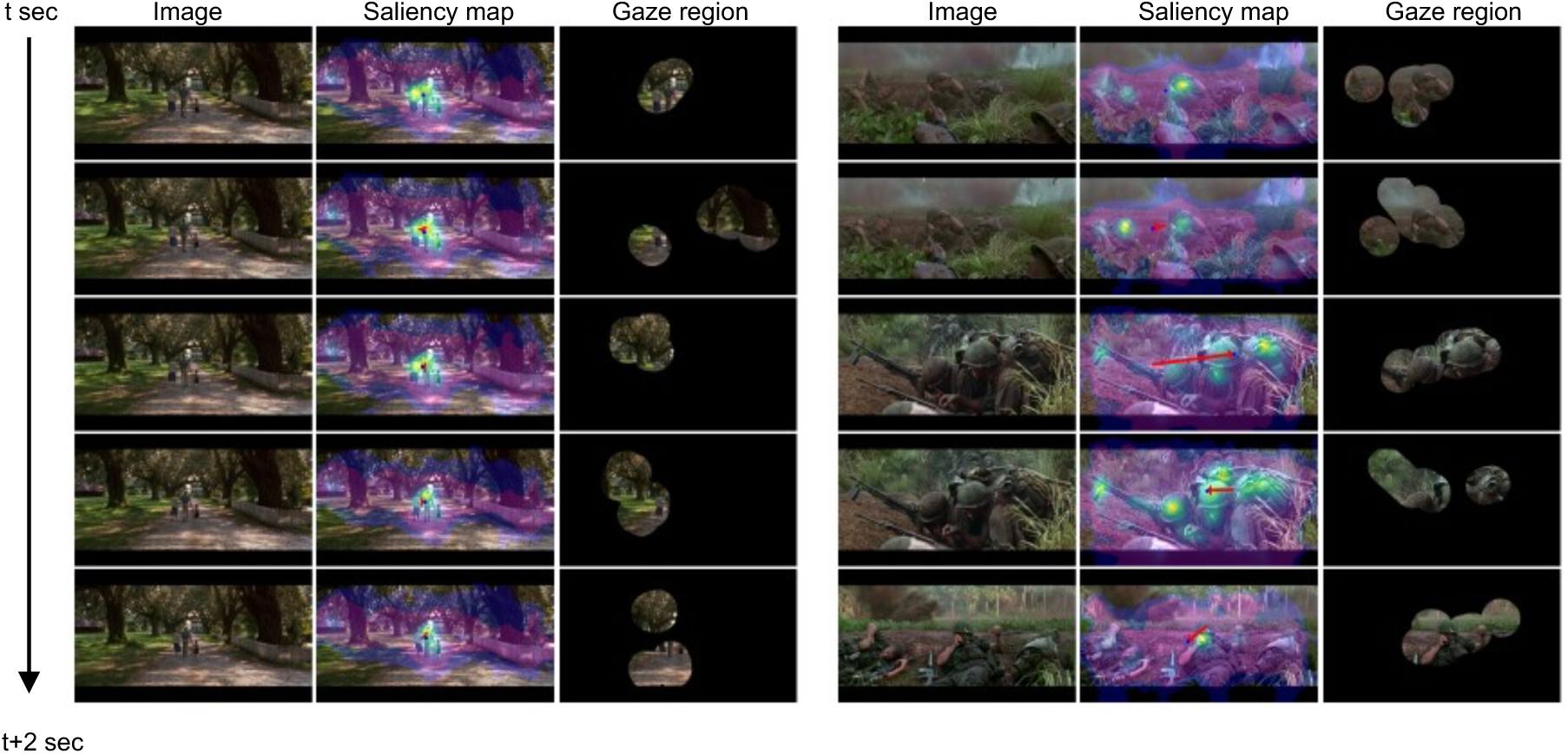
Example frames with corresponding saliency maps and gaze regions. Example frames with corresponding saliency maps predicted by DeepGaze IIE^23^ and gaze regions of Participant 01 over time. Arrows in the saliency maps indicate transitions from the previous to the current saliency centroid. When the shift of the saliency centroid is small (left), Participant 01 explored regions with low saliency as well as high-saliency regions. Conversely, when the shift is large (right), the gaze followed the saliency transitions.

**Supplementary Fig. 2:**
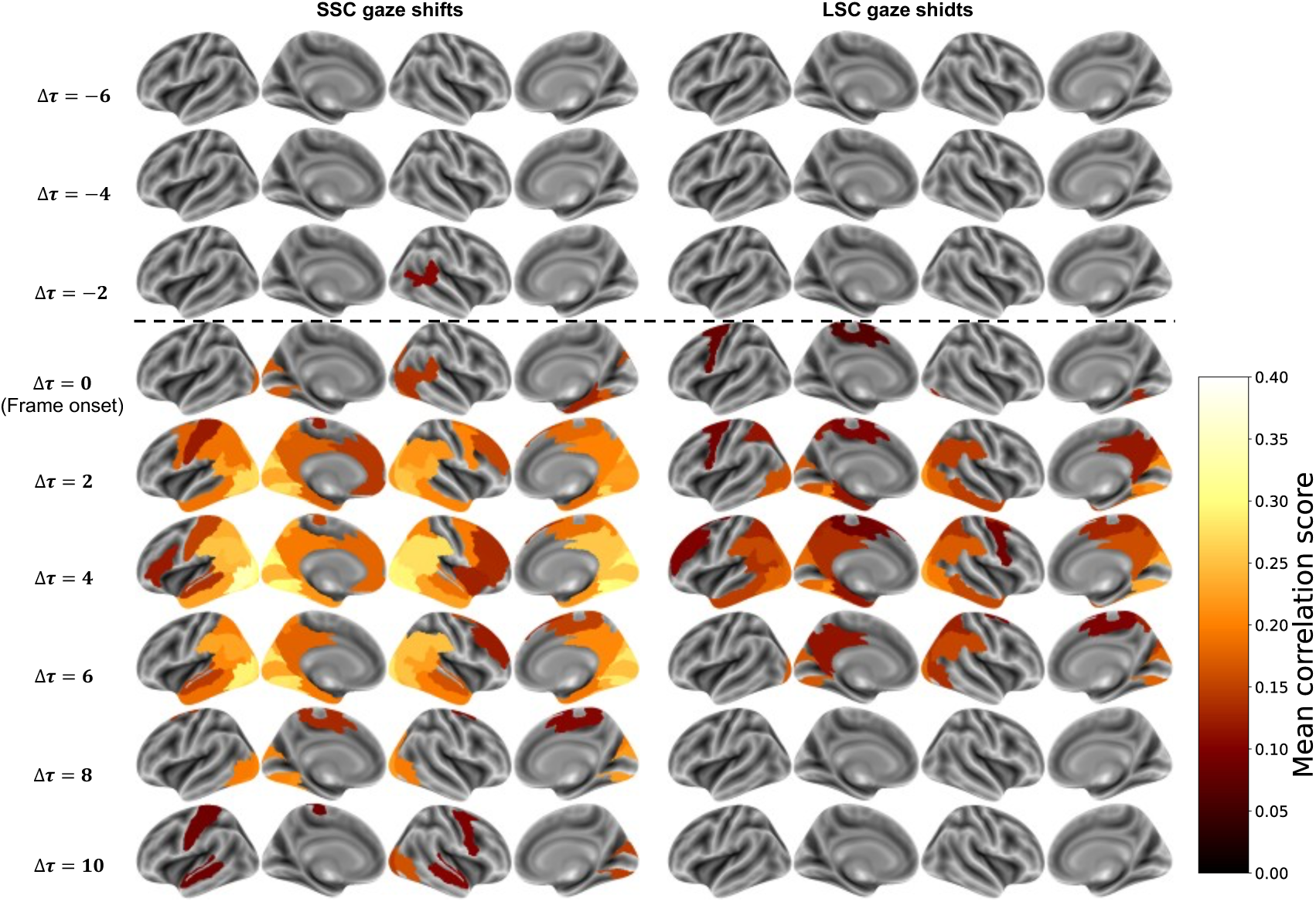
Spatiotemporal distribution of ROIs significantly predicting the visual features of gaze regions. ROIs with significant correlation scores (one-sided t-test, N = 12, Bonferroni-corrected, P < 0.05) at each Δτ are color-coded according to the mean correlation score across participants.

**Supplementary Fig. 3:**
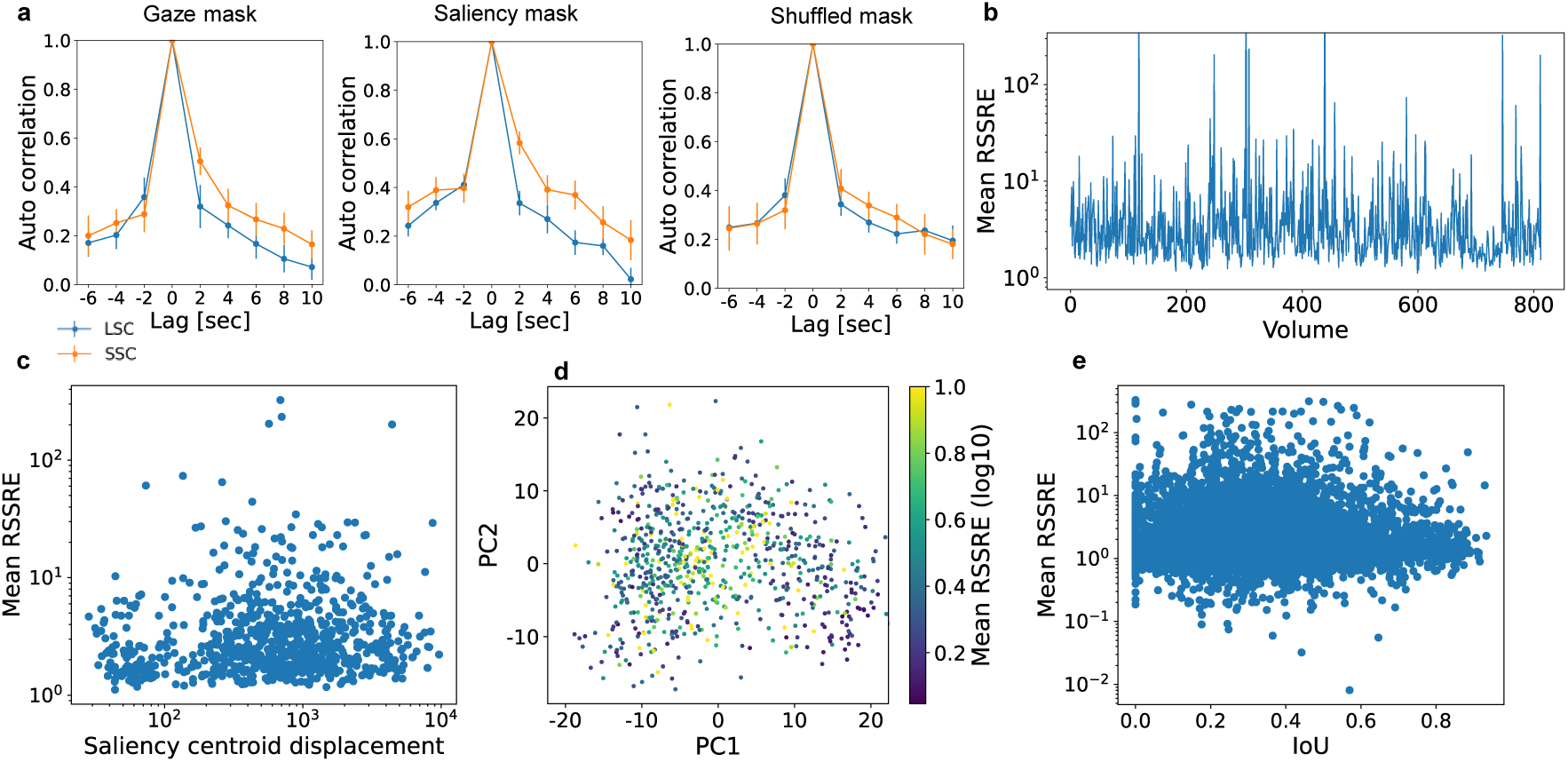
Feature-level relationships among saliency change, gaze behavior, masked image features, and the prediction error of R-TPOJ. **a**, Temporal autocorrelation of masked visual features for the gaze mask, saliency mask, and shuffled mask. There is little difference between large and small saliency change (LSC and SSC) conditions at negative lags. **b**, Temporal profile of the mean Root Sum of Squared Relative Errors (RSSRE) across participants when predicting visual features of gaze regions from BOLD responses in the right temporo-parieto-occipital junction (R-TPOJ) with a temporal lag Δ𝜏=−2. In panels **b–e**, samples with RSSRE values exceeding 3σ above the mean were excluded. **c**, Scatter plot of the logarithm of mean RSSRE across participants versus the logarithm of saliency centroid displacement. The Pearson correlation coefficient was 0.10 (*p* < 0.05). **d**, Relationship between the mean RSSRE and the distribution of visual features within the gaze region. Each dot represents a sample, and color indicates the log-scaled mean RSSRE across participants. **e**, Scatter plot of the logarithm of RSSRE and the gaze shift magnitude, quantified as the intersection-over-union (IoU) between consecutive gaze regions. The Pearson correlation coefficient was –0.017 (*p* > 0.05).

**Supplementary Fig. 4:**
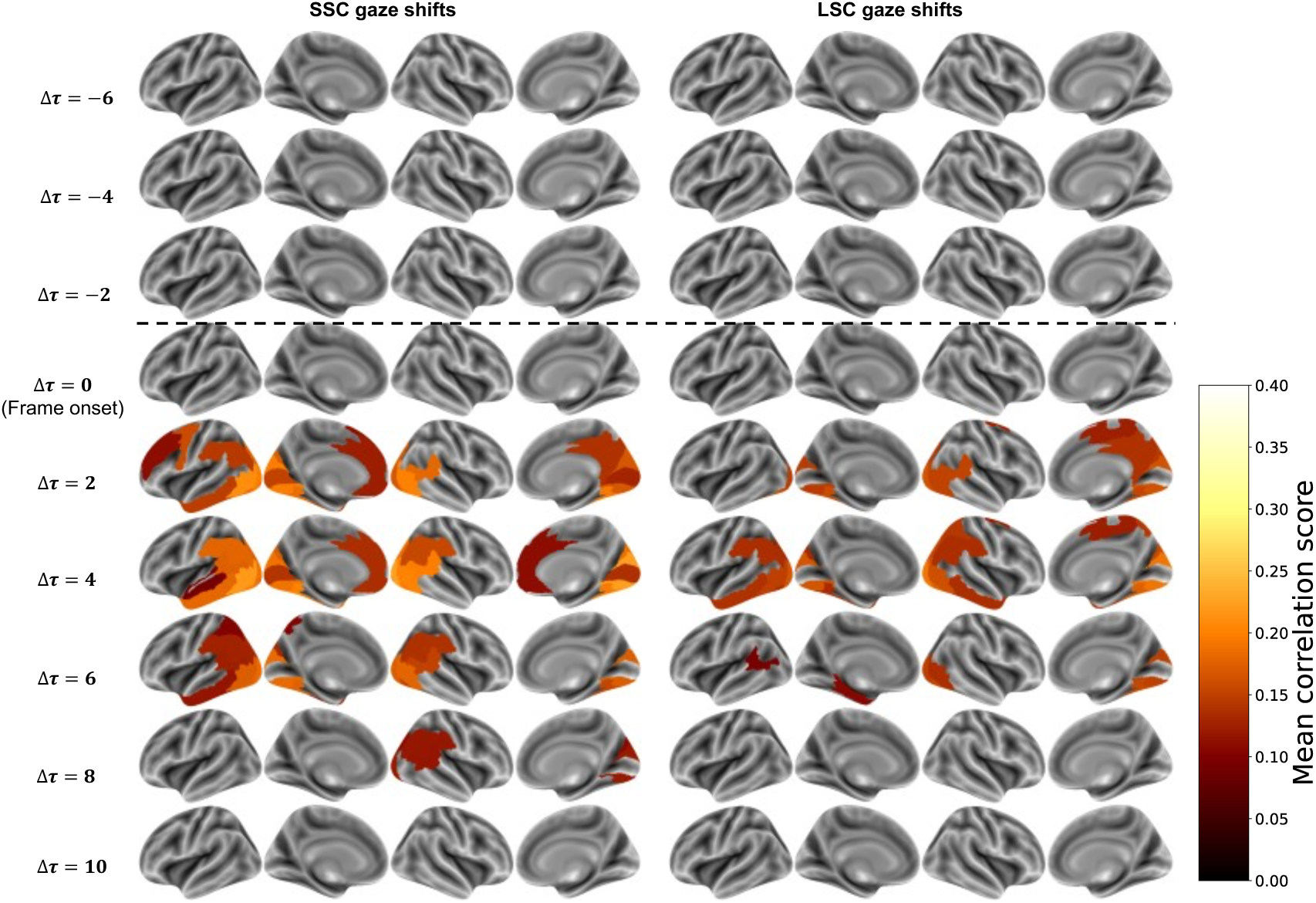
Spatiotemporal distribution of ROIs significantly predicting the visual features of saliency-masked images. ROIs with significant correlation scores (one-sided t-test, N = 12, Bonferroni-corrected, P < 0.05) at each Δτ are color-coded according to the mean correlation score across participants.

**Supplementary Fig. 5:**
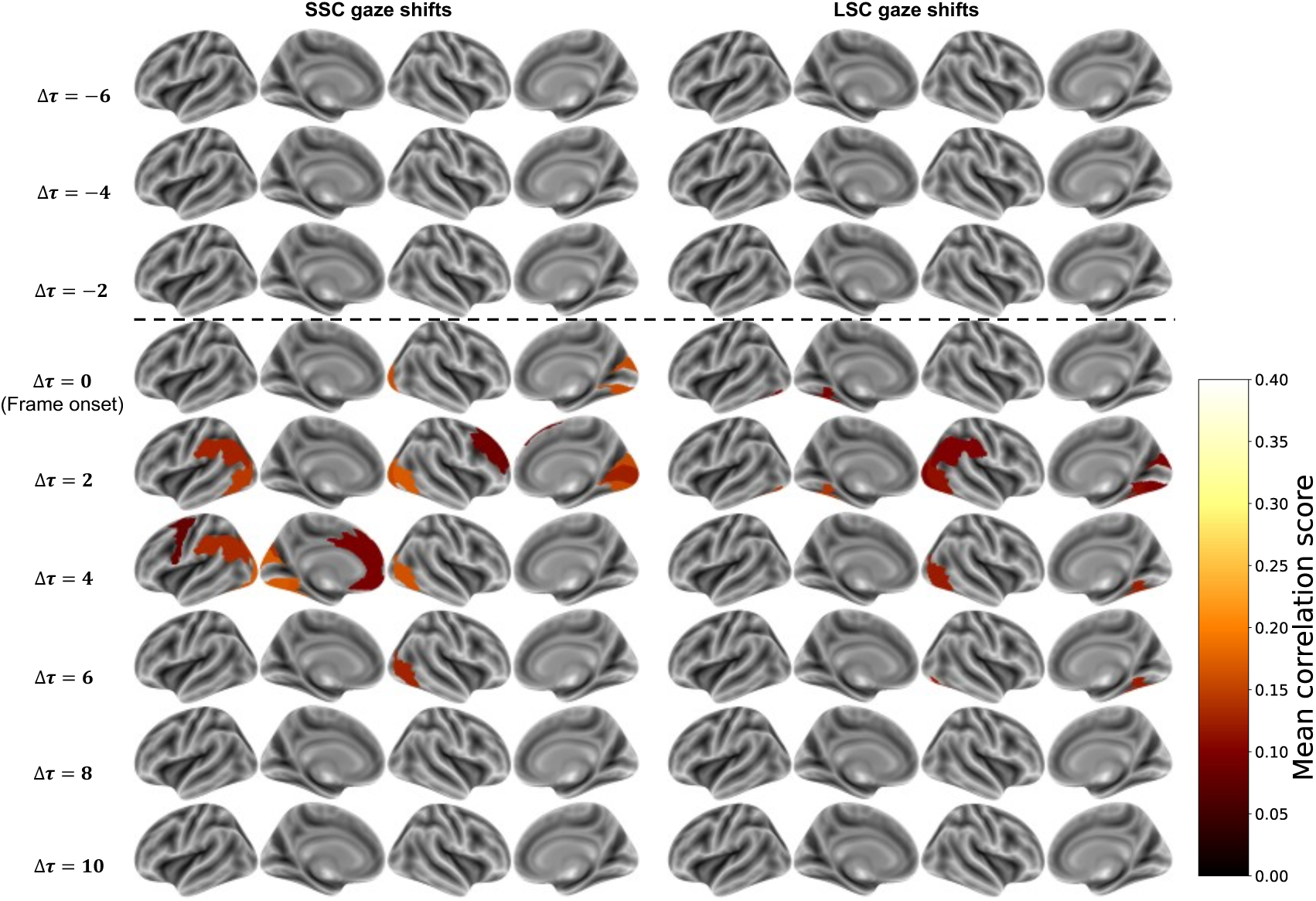
Spatiotemporal distribution of ROIs significantly predicting the visual features of shuffled-masked images. ROIs with significant correlation scores (one-sided t-test, N = 12, Bonferroni-corrected, P < 0.05) at each Δτ are color-coded according to the mean correlation score across participants.

**Supplementary Fig. 6:**
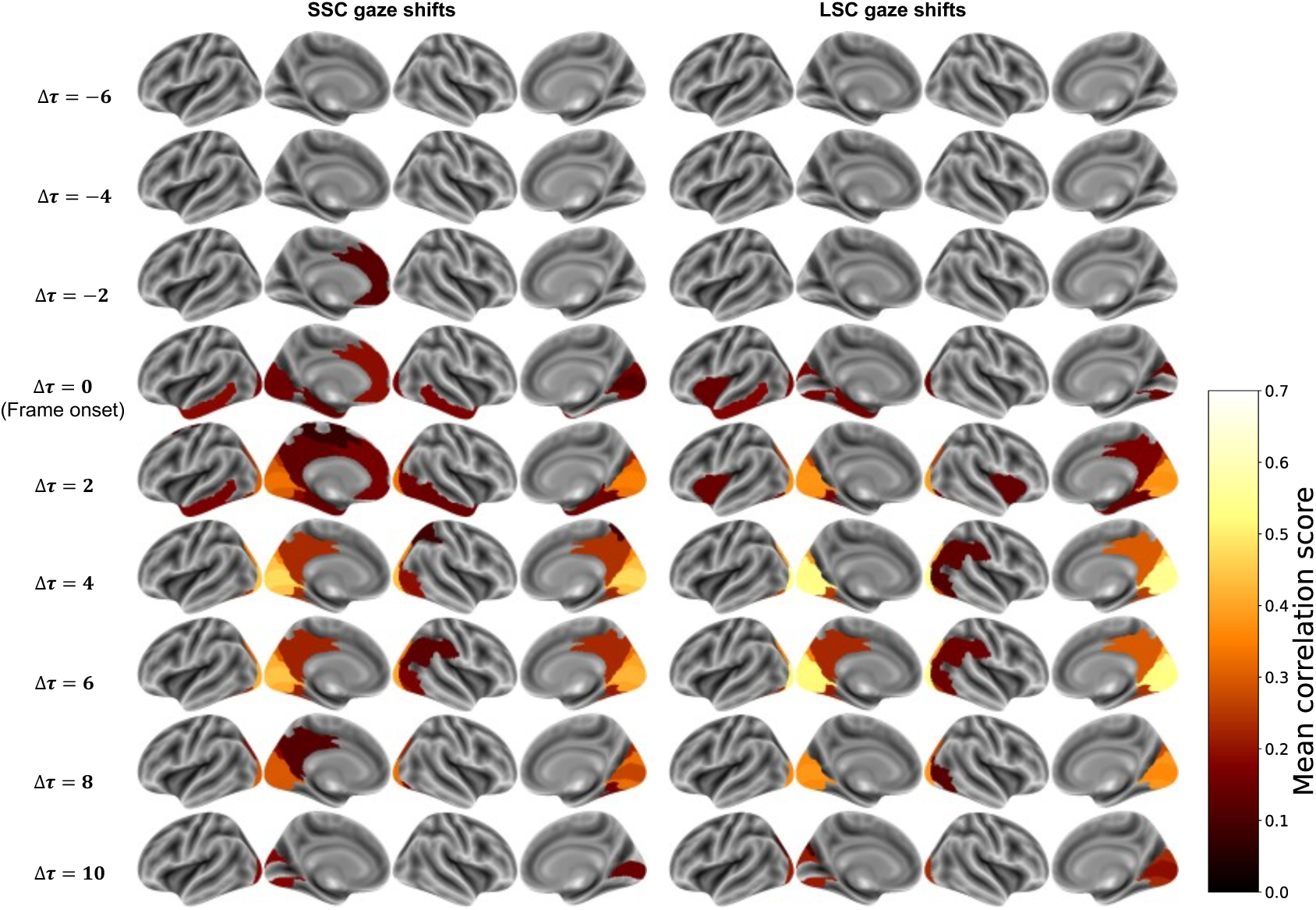
Spatiotemporal distribution of ROIs significantly predicting the gaze positions. ROIs with significant correlation scores (one-sided t-test, N = 12, Bonferroni-corrected, P < 0.05) at each Δτ are color-coded according to the mean correlation score across participants.

**Supplementary Fig. 7:**
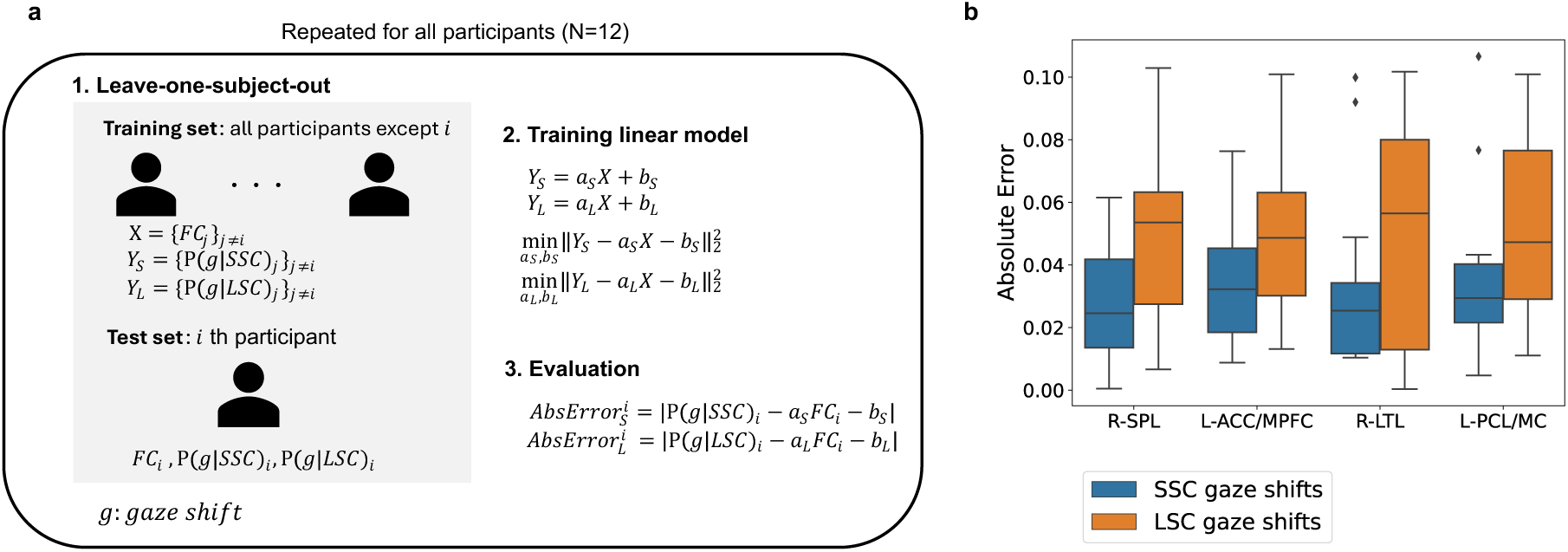
Cross-subject prediction of gaze-shift probability from functional connectivity. **a**, For each subject, a leave-one-subject-out (LOSO) analysis was performed. Pairs of functional connectivity (FC) values and probabilities of SSC or LSC gaze shifts were used to train a linear model that predicts the gaze-shift probability from FC using data from all participants except one. The model was then tested on the held-out participant to compute the absolute error between predicted and observed probability. **b**, Box plots showing absolute prediction errors for each target ROI used to compute FC. Box plots indicate the median (center line), interquartile range (box), and whiskers extending to 1.5×IQR; points beyond this range are shown as outliers. Blue and orange boxes represent SSC and LSC conditions, respectively.

## Notes

### Competing Interest Statement

The authors have declared no competing interest.

### Funding

- JST: JPMJMS2012

